# Fine scale diversification of endolithic microbial communities in the hyper-arid Atacama Desert

**DOI:** 10.1101/218446

**Authors:** Victoria Meslier, Maria Cristina Casero, Micah Dailey, Jacek Wierzchos, Carmen Ascaso, Octavio Artieda, Jocelyne DiRuggiero

## Abstract

The expansion of desertification across our planet is accelerating as the result of human activity and global climate change. In hyper-arid deserts, endolithic microbial communities colonize the rocks’ interior as a survival strategy. Yet, the composition of these communities and the drivers promoting their assembly are still poorly understood. Using a sampling strategy that minimized climate regime and biogeography effects, we analyzed the diversity and community composition of endoliths from four different lithic substrates – calcite, gypsum, ignimbrite and granite – collected in the hyper-arid zone of the Atacama Desert, Chile. By combining microscopy, mineralogy, and high throughput sequencing, we found these communities to be highly specific to their lithic substrate, although they were all dominated by the same four main phyla, *Cyanobacteria*, *Actinobacteria*, *Chloroflexi* and *Proteobacteria*. This finding indicates a fine scale diversification of the microbial reservoir driven by substrate properties. Our data suggest that the overall rock chemistry is not an essential driver of community structure and we propose that the architecture of the rock, *i.e*. the space available for colonization and its physical structure, linked to water retention capabilities, is ultimately the driver of community diversity and composition at the dry limit of life.

**Originality-Significance Statement:** In this study, we demonstrated that endolithic microbial communities are highly specific to their substrates, suggesting a fine scale diversification of the available microbial reservoir. By using an array of rock substrates from the same climatic region, we established, for the first time, that the architecture of the rock is linked to water retention and is ultimately the driver of community diversity and composition at the dry limit for life.

## INTRODUCTION

Desertification is increasing around the world as the result of human activity and climate change (Barnett *et al.*, 2005). Given that microbial communities play a critical role in desert “ecosystem services”, especially in hyper-arid deserts where plant coverage is extremely limited (Pointing and Belnap 2012; Makhalanyane *et al.*, 2015), it is therefore essential to further our understanding of the functioning of these communities and of their resilience mechanisms to climate and anthropomorphic perturbations. Endolithic microbial communities (inhabiting rocks’ interior) are well represented in hot, cold and polar deserts around the world (Walker and Pace 2007; Omelon 2008; Wierzchos *et al.*, 2012a). These communities are relatively simple and, as such, provide good model systems to elucidate adaptive mechanisms to the dry limit of life (Walker and Pace 2007). Often considered as the last refuge for life in extreme deserts, the endolithic habitat provides protection against harmful solar irradiance, enhances the water budget by up-taking and retaining water, buffers high thermal fluctuations, and provides physical stability to microbial communities (Friedmann 1980; Walker and Pace 2007; Wierzchos *et al.*, 2012a). The capacity of the rock substrate to harbor life, *i.e.* its bioreceptivity, is determined by its chemical and physical properties (Wierzchos *et al.*, 2012a). Depending on the structure of the lithic substrate, the colonization zone can be found inside pores beneath the rock surface (cryptoendolithic), in cracks and fissures (chasmoendolithic), or inside pores in the bottom part of the rock (hypoendolithic) (Golubic *et al.*, 1981; Wierzchos *et al.*, 2011).

In hyper-arid deserts, endolithic microbial communities have been found to colonize a number of rock substrates, including carbonates (de los Ríos *et al.*, 2004; Horath *et al.*, 2006; Tang *et al.*, 2012; DiRuggiero *et al.*, 2013; Crits-Christoph *et al.*, 2016; Tang *et al.*, 2016; Kidron and Temina 2017), gypsum (Dong *et al.*, 2007; Ziolkowski *et al.*, 2013; Wierzchos *et al.*, 2015), gypsum crust (Hughes and Lawley 2003; Stivaletta *et al.*, 2010; Wierzchos *et al.*, 2011; Canfora *et al.*, 2016), halite (Wierzchos *et al.*, 2006; Davila *et al.*, 2008; de los Ríos *et al.*, 2010; Robinson *et al.*, 2015), ignimbrite (Wierzchos *et al.*, 2013; Cámara *et al.*, 2014), granite (Friedmann and Kibler 1980; de los Ríos *et al.*, 2005; de los Ríos *et al.*, 2007; Li *et al.*, 2013; Büdel *et al.*, 2008), and sandstone (McKay and Friedmann 1985; Wierzchos and Ascaso 2001; Ascaso and Wierzchos 2002; de los Ríos *et al.*, 2004; Pointing *et al.*, 2009; Archer *et al.*, 2017). The great variety of rocks supporting endolithic life shows the versatility of microorganisms to colonize a wide array of substrates and their incredible potential for adaptation to extreme environmental conditions.

Characterization of the structure and composition of these communities has shown that endoliths all harbor phototrophic primary producers, co-occurring with a wide range of heterotrophic consumers, and seeded from a “metacommunity”, *i.e.* a microbial reservoir available for colonization (Walker and Pace 2007; Wierzchos *et al.*, 2012a). Given the extreme environmental context, several studies have shown that water is the most important factor for survival of endolithic communities in hyper-arid deserts. Water can originate from sparse precipitations but also from dew, fog, deliquescence, capillary condensation, or snow melt in polar deserts and, its availability is dependent on substrate properties (Friedmann *et al.*, 1988; Omelon *et al.*, 2006; Büdel *et al.*, 2008; Davila *et al.*, 2008; Wierzchos *et al.*, 2012b; Robinson *et al.*, 2015). However, the relationship between substrate properties and the diversity of the communities inhabiting those substrates remains poorly understood.

Although culture-independent methods combined with microscopy techniques have led to a more complete understanding of endolithic communities and their habitat, a broader picture of the endolithic community structures and compositions is still lacking. Indeed, most of the studies have focused on one or two rock substrates at a time, making direct comparisons across substrates difficult, if not impossible at the molecular level. In this study, we conducted a large-scale analysis of the composition of 47 endolithic microbial communities found in calcite, gypsum, ignimbrite and granite rocks collected in the same climate regime of the Atacama Desert. We combined field measurements, microscopy, geochemical characterization, and 16S rDNA amplicon high-throughput sequencing to address the communities’ specific features and identify the drivers promoting their assembly. Within the same climate regime, we found endolithic communities highly specific to their substrates, suggesting a fine scale diversification of the available microbial reservoir.

## RESULTS

We combined microclimate measurements, mineralogy and microscopy analyses, and high throughput culture-independent molecular data to identify the factors underlying the structure and composition of microbial assemblages in endoliths from the hyper arid Atacama Desert. Illumina sequencing of 47 samples resulted in a total of 1,773,072 V3-V4 SSU rDNA reads, with an average number of paired-end reads per sample of 17,469 +/- 4,375 and an average length of 452 +/- 11 bp, providing an unprecedented level of detail for these communities. Slopes of the rarefaction curves indicated that most of the community was sampled at the 3,000-sequence level (**Supplementary Information S4**).

### Microclimate data

Forty-seven samples from 4 different rock substrates were collected from two sampling sites, Valle de la Luna area (VL) and Monturaqui area (MTQ), located in the hyper-arid zone of the Atacama Desert (**Figure 1** and **Supplementary information S1**). Climate data were recorded over a period of at least 22 months, showing similar climate regimes for both sites (**Figure 1**). The mean air temperature was about 16°C, with strong amplitude between minima and maxima (from -4°C to 49°C). The average diurnal PAR was ~ 1000 µmol photons m^-2^ s^-1^ with a maximum of 2553.7 µmol photons m^-2^ s^-1^, substantiating the extremely intense solar irradiance found in this region (Cordero *et al.*, 2014). Both sites experienced extremely dry conditions, with an average air relative humidity (RH) of about 20% with frequently lows of ~5% and 1% in VL and MTQ, respectively. Precipitations were extremely scarce with mean annual values of 24.5 and 27.8 mm/y in MTQ and VL, respectively.

**Figure 1:**
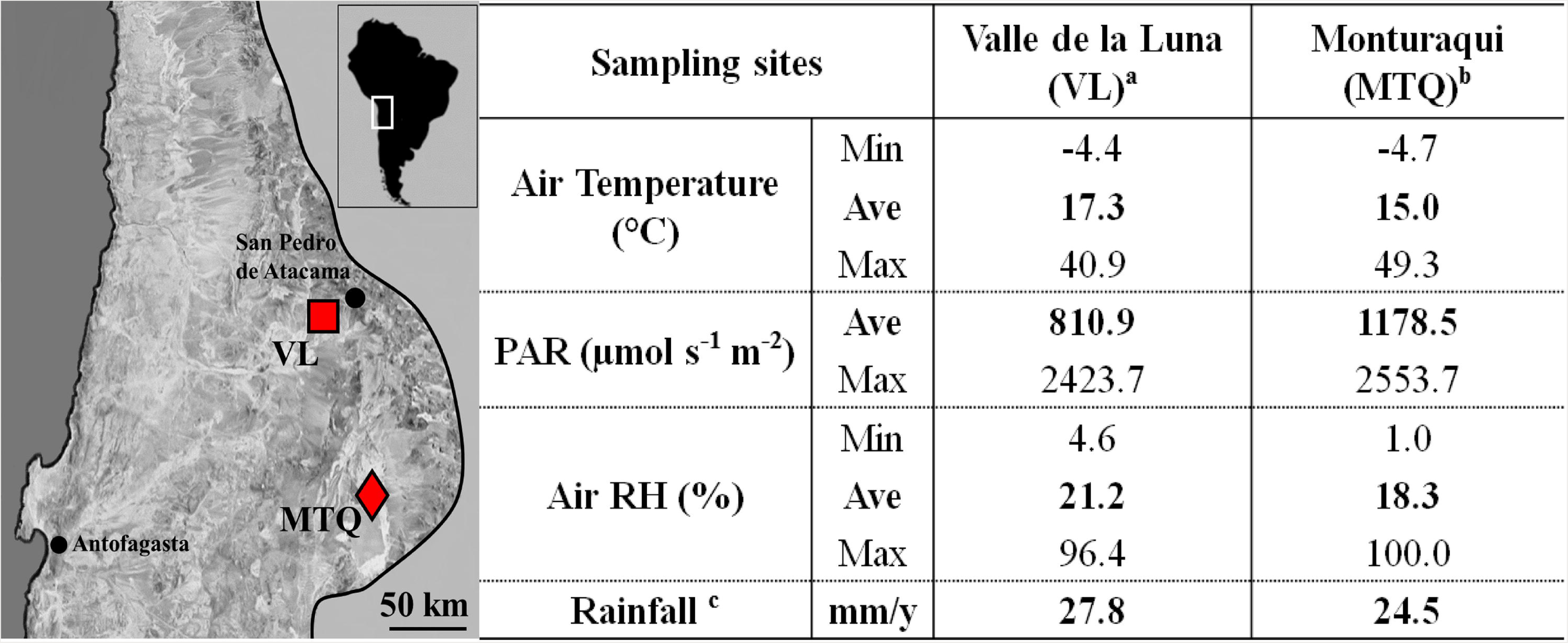
**Sampling locations in the Atacama Desert and microclimate data.** Map of the Atacama Desert in Chile, with sampling locations: VL, Valle de la Luna (red square) and MTQ, Monturaqui area (red diamond). ^a^data recorded from April 2013 to December 2015 (32 months - this paper); ^b^data from January 2011 to February 2013 (22 months) (Wierzchos *et al.*, 2015); cdata extracted from DiRuggiero *et al.* (2013).

### Mineralogy of the substrates

X-ray diffraction analysis (XRD) showed that the samples collected in VL were sedimentary carbonate rocks composed of calcite (**Table 1**). In the MTQ area, the gypsum rocks were mainly composed of calcium sulfate with minor minerals such as cristobalite, calcite and potassium feldspar. These evaporitic rocks also contained the mineral sepiolite. In contrast, the mineral composition of the ignimbrite and granite rocks was more complex with the pyroclastic ignimbrite rocks composed of andesine, disorded sodian anorthite, and biotite. The granite rocks were composed of quartz, potassium feldspar with small amounts of plagioclase and biotite. The water soluble ions measured for each of the substrate type were remarkably low, regardless of the chemical complexity of the substrate, and often close to the detection limits of the ICP and IC methods (**Supplementary Information S3**).

**Table 1:** Mineralogical composition of rock substrates obtained by XRD

| Substrate | XRD analysis | Chemical formula | References |
| --- | --- | --- | --- |
| **Calcite** | Calcite (100%) | $\text{CaCO}_3$ (100%), | DiRuggiero *et al.*, 2013 |
| **Gypsum** | Gypsum (96–98%), cristobalite, calcite, potassium feldspar (2-4%) | $\text{CaSO}_4 \cdot 2\text{H}_2\text{O}$ (96-98%), $\text{SiO}_2$, $\text{CaCO}_3$, $\text{KAlSi}_3\text{O}_8$ | Wierzechos *et al.*, 2015 |
| **Ignimbrite** | Andesine (51%), disordered sodian anorthite (42%), biotite (7%) | $(\text{Ca}, \text{Na})(\text{Al}, \text{Si})_4\text{O}_8$ (51%), $\text{CaAl}_2\text{Si}_2\text{O}_8$ (42%), $\text{K}(\text{Mg}, \text{Fe})_3\text{AlSi}_3\text{O}_{10}(\text{OH})_2$ (7%) | Wierzechos *et al.*, 2013 |
| **Granite** | Quartz (25%) potassium feldspar (22%) plagioclase (34%), biotite (16%) other (3%) | $\text{SiO}_2$ (25%), $\text{KAlSi}_3\text{O}_8$ (22%), $\text{K}(\text{Mg}, \text{Fe})_3\text{AlSi}_3\text{O}_{10}(\text{OH})_2$ (6%), other (3%) | this study |

### Endolithic colonization zones

Cross sections of the rocks revealed a significant heterogeneity in the micromorphology and structure of the lithic substrates (**Figure 2 A1-D1, Table 2**). Calcite rocks, composed of laminated layers covered by a hard surface layer with microrills (DiRuggiero *et al.*, 2013), displayed large irregular fissures and cracks perpendicular or parallel to the rock surface, with a green colonization zone as deep as 1.5 cm (**Figure 2-A1, Table 2**). Microbial cells were organized in large clusters along the cracks and fissures of this substrate and adhered to different minerals without preference (**Figure 2-A2**). SEM-BSE revealed dense arrangements of cyanobacterial cells embedded in concentric sheets of extracellular polymeric substance (EPS) that were filled by heterotrophic bacteria (**Figure 2-A3**). The colonization zone within gypsum was mostly cryptoendolithic, close to the compact gypsum surface layer, and two characteristic pigment layers, orange for carotenoids and green for chlorophylls, were organized with depth (Wierzchos *et al.*, 2015, Vitek *et al.*, 2016) (**Figure 2-B1, Table 2**). *Cyanobacteria* were found among lenticular gypsum crystals, filling up vertically-elongated pores, and aggregated around sepiolite nodules (**Figure 2-B2**). This clay mineral, with high water retention capacity, was previously identified in gypsum by Wierzchos *et al.* (2015). Detailed SEM-BSE image of the gypsum (**Figure 2-B3)** showed *Cyanobacteria* with different micromorphology (larger cells) accompanied by heterotrophic bacteria. Ignimbrite rocks had a consistent and narrow cryptoendolithic colonization zone, characterized by a vitrified foam-like structure under a brown-colored varnish covering the rock surface (**Figure 2-C1, Table 2**) (Wierzchos *et al.*, 2013). Microscopically, this substrate showed small elongated vesicles among a glass shards matrix. Cells filled these narrow spaces, while others, not connected to the surface, showed no signs of microbial colonization **(Figure 2-C2)**. Detailed SEM-BSE view revealed dense cyanobacterial aggregates and heterotrophic bacteria attached to the glass shards **(Figure 2-C3)**. Granite rocks, devoid of a surface crust, had a dark green chasmoendolithic colonization within random arranged fissures and cracks under the rock surface **(Figure 2-D1)**. Dense microbial aggregates were found among grains of quartz and feldspars **(Figure 2-D2)**. A detailed view showed cyanobacterial cells associated with rod shape heterotrophic bacteria **(Figure 2-D3)**.

**Table 2:**
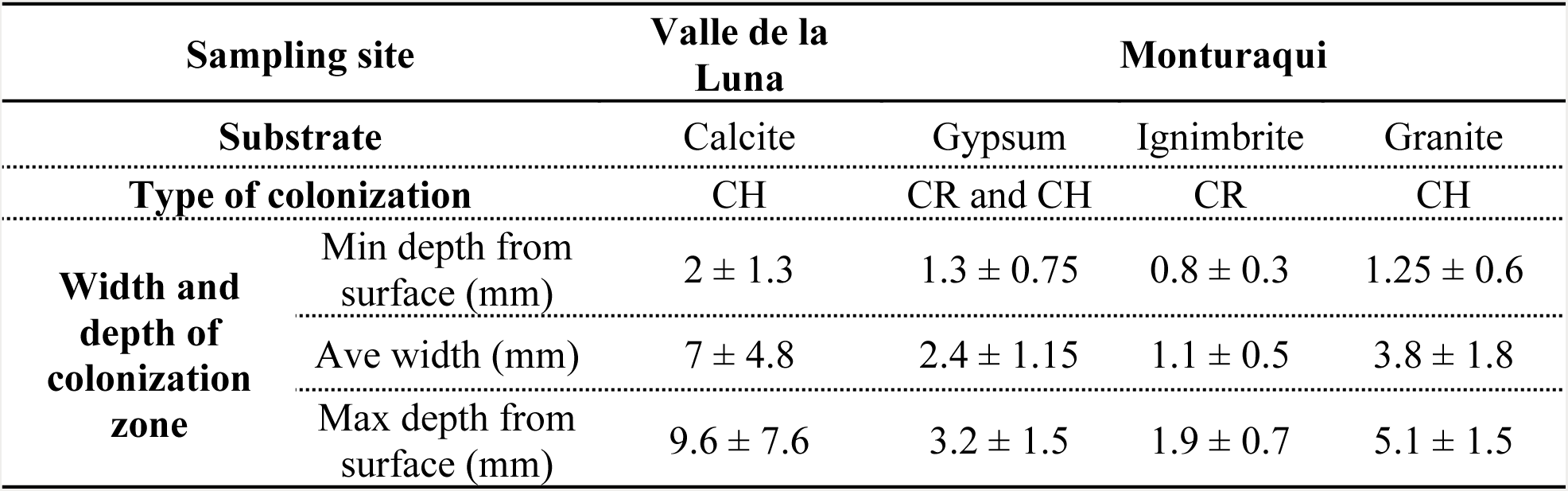
Characteristics of endolithic colonization zones.

Reported for each substrate are the type of colonization (CR: cryptoendolithic, CH: chasmoendolithic), width, minimum and maximum depth of colonization in mm (minimum and maximum depths were measured from top surface). Values are the results of 3 measurements per individual rock for at least 4 rocks per substrate.

**Figure 2:**
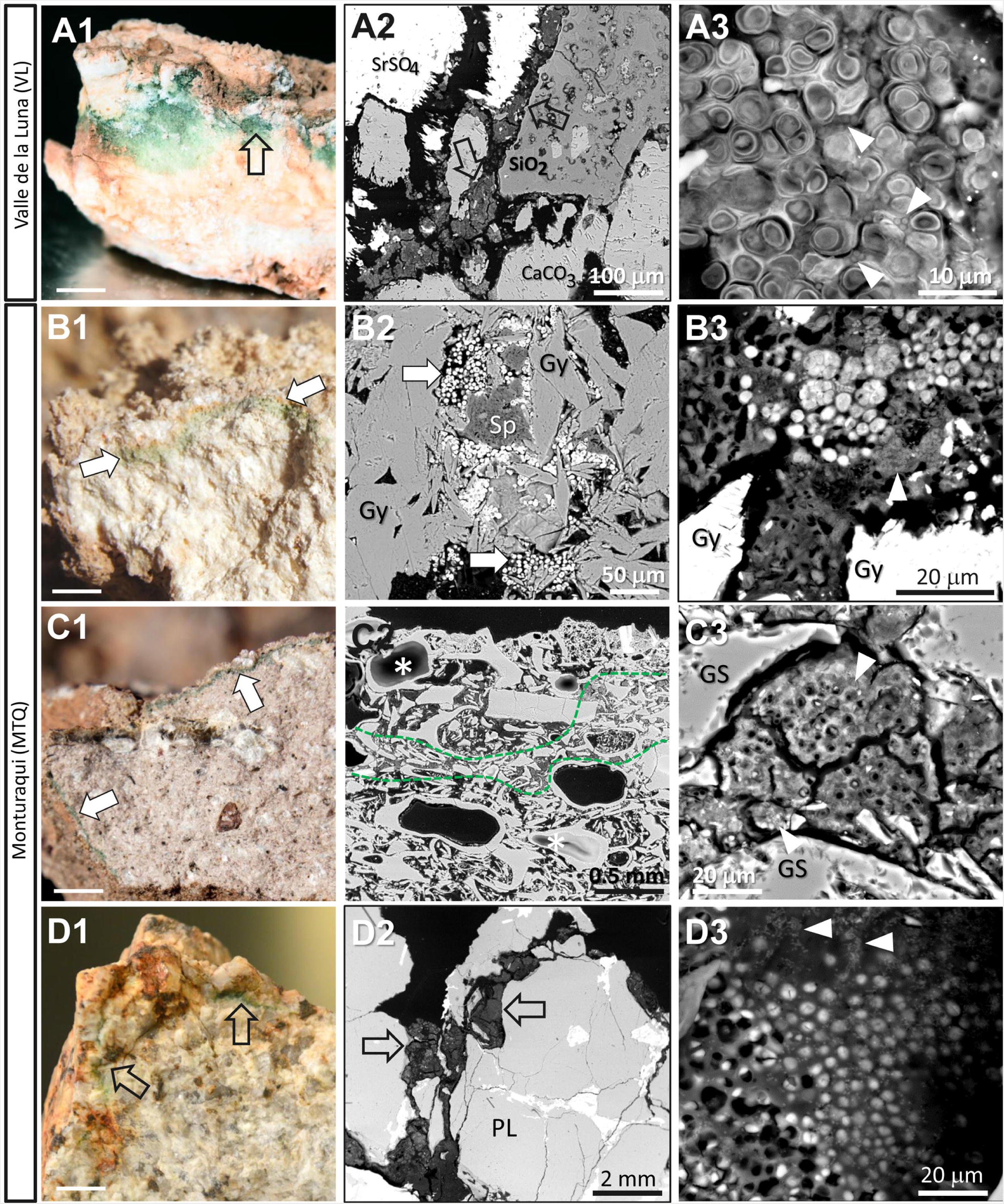
**Cross-sections and SEM-BSE images of endolithic colonization zones for calcite (A), gypsum (B), ignimbrite (C) and granite (D)**. (**A1-D1**) cross-sections of colonized rocks with variation in depth, width, distribution, and color of the endolithic habitats; open black and filled white arrows indicates chasmoendolithic **(A1 and D1)** and cryptoendolithic **(B1 and C1)** habitats, respectively; scale bars: 1 cm. Series **A2-D2 and A3-D3** shows SEM-BSE images revealing aggregates of endolithic communities inside the cracks and pores of the lithic substrates; (**A2-A3)** fissures within calcite filled by chasmoendoliths between celestine (SrSO_4_), silica (SiO_2_), and calcite (CaCO_3_) minerals; (**B2-B3)** aggregates of cyanobacteria among gypsum (Gy) crystals and surrounding sepiolite (S) nodules; (**C2-C3)** aggregates of cyanobacteria within bottle shape pores of ignimbrite, green lines represent the cryptoendolithic colonization zone; and (**D2-D3)** dense cyanobacterial cell aggregates within cracks of granite. White arrowheads indicated heterotrophic bacteria associated to cyanobacterial aggregates.

### Structure and composition of endolithic communities

High throughput sequencing of 16S rDNA amplicons was used to characterize communities across 47 samples and 4 substrates. Diversity metrics, calculated from OTUs clustered at 97%, revealed two significantly different groups of substrates with high and low diversity (T-statistic = 10.8, p-value < 0.0001) (**Supplementary Information S4**). The granite and ignimbrite substrates grouped at the lower end of diversity (<220 OTUs; Shannon index <5) with no significant differences in alpha diversity indices (Observed Richness; T-statistic=1.52, p-value=0.17). In contrast, the calcite and gypsum substrates harbored high diversity (> 400 OTUs; Shannon index ~ 6) and significant differences in their observed richness (T-statistic=2.57, p-value=0.016). It is notable that the gypsum and ignimbrite substrates, collected side by side in the field (photo in **Supplementary Information S1**), had the highest (545) and lowest (126) number of OTUs, respectively.

A total of 14 bacterial phyla, but no *Archaea*, were found across all substrates and distinct high-ranked taxon dominated all substrates. *Cyanobacteria*, *Actinobacteria*, *Chloroflexi* and *Proteobacteria* were the most abundant phyla, representing 86 to 98% of the total community (**Figure 3-A**). *Cyanobacteria* dominated the communities in all substrates (**Figure 3-B**) with relative abundance from ~60% to more than 70%. *Actinobacteria* was the second most abundant phylum but never exceeded 30% of the total community composition (**Figure 3-C**). One-way ANOVA confirmed significant differences of the mean relative abundance for the 4 main phyla across substrates (**Figure 3** and **Supplementary Information S5**). *Deinococcus*, *Gemmatimonadetes, Bacteroidetes* and *Armatimonadetes* were found in all substrates at relatively low abundance, between 1 and 5%, with higher levels in calcite and gypsum. Six phyla, *Acidobacteria*, *Saccharibacteria*, *Nitrospirae*, *Planctomyces*, *Verrucomicrobia* and candidatus WHCB1-60, were found at very low abundance, ≤ 1%, in calcite and gypsum and sporadically detected in granite and ignimbrite (**Supplementary Information S6**). Strong anti-correlations of the relative abundance of *Cyanobacteria* were found with that of *Actinobacteria* and *Proteobacteria* (rho_spearman_ = -0.821 and -0.850, respectively, p-values < 1.10^−4^) and *Chloroflexi* (rho_spearman_ = -0.459, p-value = 0.001) for all substrates (**Figure 3** and **Supplementary Information S5**). Diversity metrics were also strongly anti-correlated with the abundance of *Cyanobacteria* (rho_spearman_ =-0.718 and -0.833 for observed richness and Shannon indexes, respectively, p-values < 1.10^−4^), while these metrics were positively correlated to the abundance of the other 3 main phyla (rho_spearman_ = from 0.308 to 0.758, p-values from 1.10^−4^ to 0.03).

**Figure 3:**
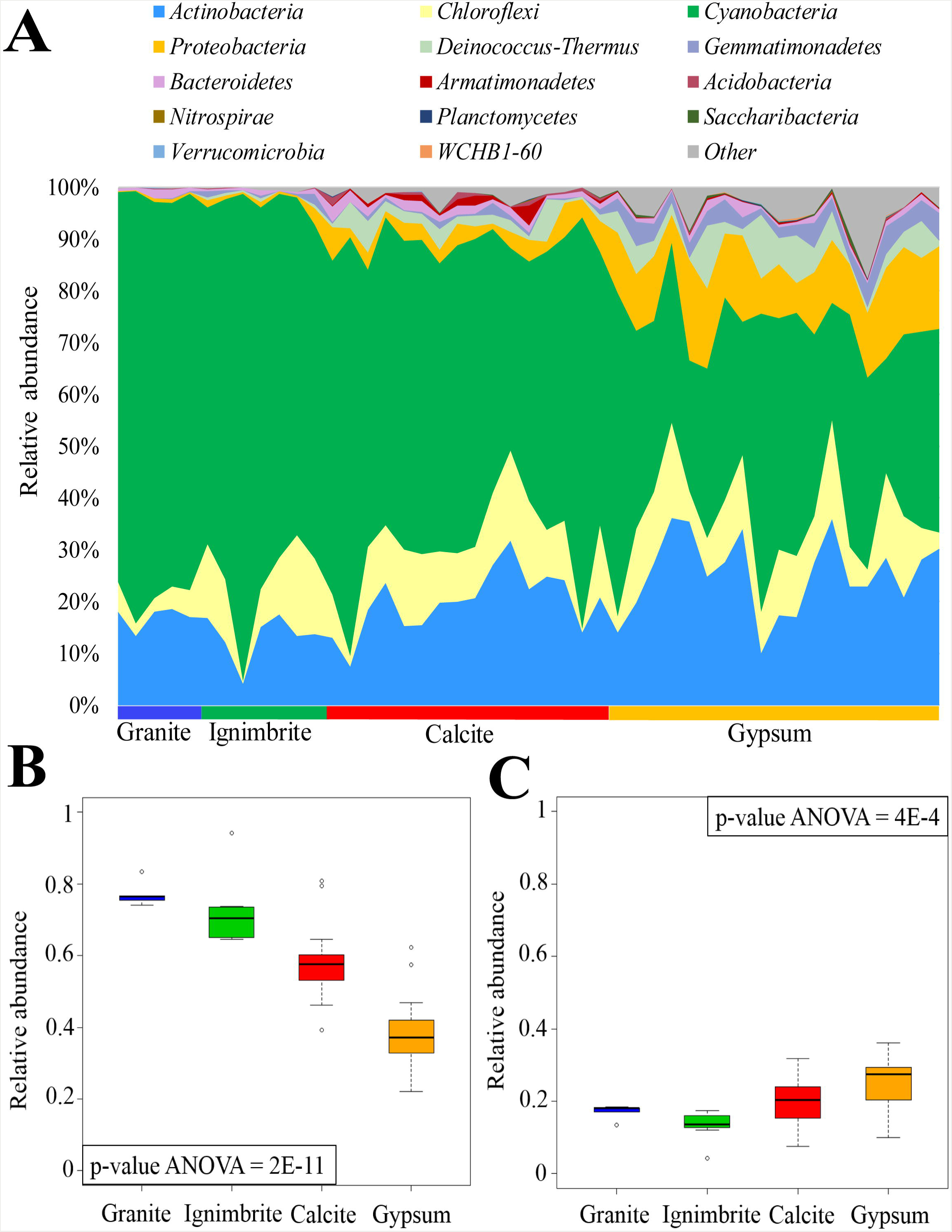
**Composition of endolithic microbial communities.** A total of 14 phyla were found using the V3-V4 region of the 16S rRNA gene with OTUs clustering at 97%. (**A**) relative abundance (%) of phyla detected across all samples, others indicate unassigned reads. Boxplots of relative abundance for *Cyanobacteria* (**B**) and *Actinobacteria* (**C**) per substrate-type; one-way ANOVA p-values are reported for each boxplot; granite is in blue, ignimbrite in green, calcite in red, and gypsum in orange.

### Partitioning of diversity

We further examined the endoliths’ community composition at a higher level of resolution using Beta-diversity analyses. A hierarchical clustering analysis, based on pairwise Bray-Curtis dissimilarity indices, was used to build an UPGMA tree (**Figure 4**). This analysis revealed that samples clustered into 4 distinct groups and according to their substrate type. The ignimbrite and granite samples were the least distant from each other, which was consistent with their low observed richness, whereas gypsum samples were as distant to the ignimbrite/granite samples as the calcite samples were, regardless of their proximity in the field (**Figure 4**). This substrate-type grouping was supported by PCoA plots generated from unweighted and weighted UniFrac distance matrices (**Supplementary Information S7)**. Adonis and ANOSIM tests, performed with substrate categories, confirmed the statistical significance of the grouping (R^2^ = 0.61 and R=0.99 with p-values = 0.001 for adonis and ANOSIM, respectively).

**Figure 4:**
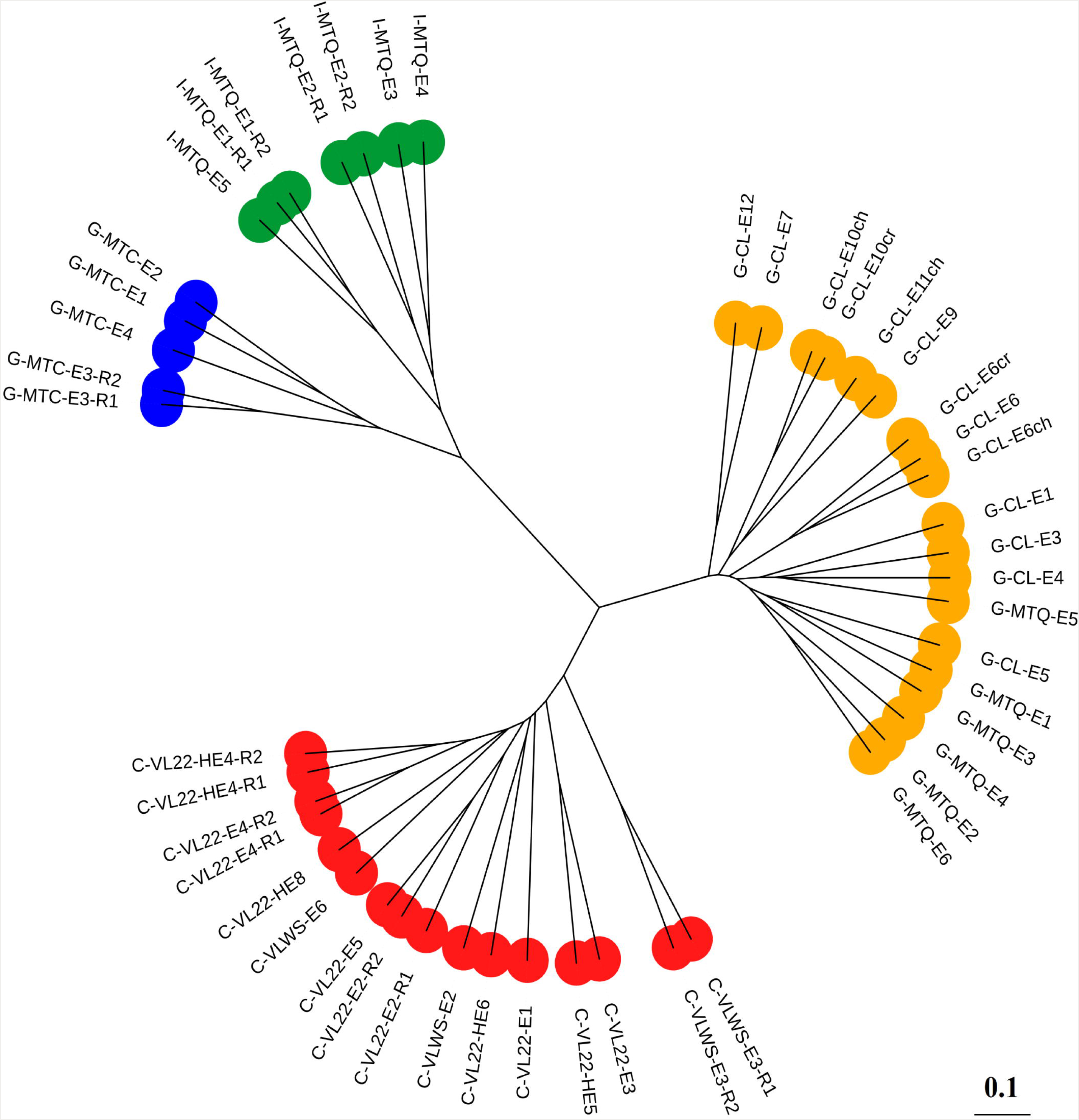
**Clustering of endolithic communities by substrate-type.** Unweighted Pair Group Method with Arithmetic Mean (UPGMA) dendrogram was constructed using Bray-Curtis dissimilarity indices and visualized using iTOL. Granite in blue, ignimbrite in green, calcite in red, and gypsum in orange. Scale bar indicates 10% dissimilarity between samples.

Despite *Cyanobacteria* being the most abundant phyla across all substrates, it was less diverse phylogenetically than *Actinobacteria* (161 versus 426 OTUs). A fine-scale analysis of the 30% most abundant *Cyanobacteria* and *Actinobacteria* OTUs (50 and 129 OTUs, respectively), comparing their relative abundance across samples, revealed clusters of OTUs preferentially associated with substrate types (**Figure 5A** and **Supplementary Information S8**). To test the statistical basis of this observation, we performed a differential abundance analysis using DESeq2 between the high (gypsum and calcite) and low (ignimbrite and granite) diversity groupings defined above by alpha diversity metrics. All *Cyanobacteria* OTUs, except for 2, were significantly assigned to either the low (n=23) or the high diversity groups (n=25) (adjusted p-values <0.05; **Figure 5B),** supporting the clusters visualized in **Figure 5A**. Similarly, top *Actinobacteria* OTUs were significantly assigned to either the low (n=14) or the high (n=86) diversity groupings, whereas 29 of them could not be significantly assigned (**Supplementary Information S8**).

**Figure 5:**
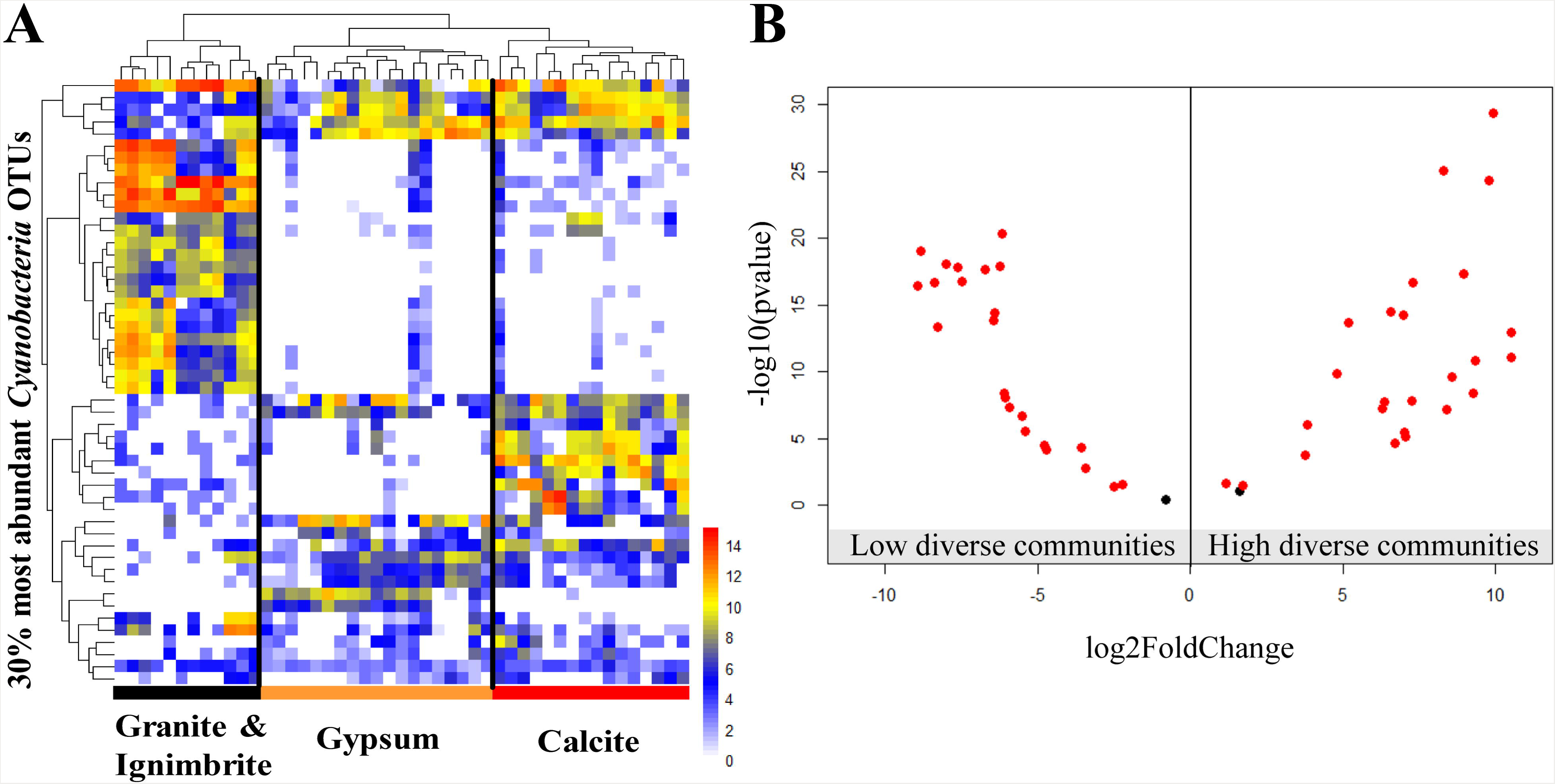
**Differential abundance of the 30% most abundant *Cyanobacteria* OTUs (n=50)**. (**A**) Heatmap of *Cyanobacteria* OTUs clustered by sample (columns) and relative abundance (rows). Color scales refer to the normalized abundance of each OTU across samples (log-transformed for visualization purposes). Granite and ignimbrite samples are in black, calcite in red, and gypsum in orange. (**B**) Volcano plots of *Cyanobacteria* OTUs in low (granite and ignimbrite) and high (gypsum and calcite) diversity endolithic communities. The log_2_ fold change, calculated for each OTU using DESEq2, was plotted against the adjusted negative log10 p-values (red; padj [adjusted p-values] <0.05, black; padj = 0.05).

A phylogenetic analysis of the major *Cyanobacteria* OTUs showed that most of them were assigned to *Chrooccocidiopsis* (~70% of the OTUs) (**Supplementary Information S9**). *Actinobacteria* OTUs were mainly identified as members of the *Solirubrobacter, Rubrobacter, Actinobacterium, Euzebya, Microlunatus, Molestobacter* and *Geodermatophilus* genus (**Supplementary Information S9**). Both maximum likelihood trees showed that substrate-specific clusters of OTUs, in the low and high diversity communities, had members belonging to similar taxonomic groups.

## DISCUSSION

In this study, we addressed the drivers promoting the assembly of endolithic microbial communities in different substrates all collected in the same hyper-arid climate regime of the Atacama Desert. The rocks were collected within a small geographic area, sometime side by side in the field, in an effort to minimize any effect from biogeography. While previous studies have demonstrated the impact of climate regime, in particular the availability of atmospheric water, in shaping the structure of lithic communities (Friedmann *et al.*, 1988; Cockell *et al.*, 2002; Pointing *et al.*, 2007; Büdel *et al.*, 2008; Davila *et al.*, 2008; Stomeo *et al.*, 2013; Robinson *et al.*, 2015), most have only focused on one type of rock substrate or reported limited data on community diversity. For example, the diversity of microbial communities in calcite rocks (DiRuggiero *et al.*, 2013) and halite nodules (Robinson *et al.*, 2015) from the Atacama Desert was found to be strongly correlated with atmospheric moisture. Similarly, shifts in community diversity and structure were reported for quartz hypoliths along moisture transects in China and Namib deserts (Pointing *et al.*, 2007; Stomeo *et al.*, 2013). However, to our knowledge, no study has compared community assembly over a wide range of endolithic substrates and without the confounding effect of different climate regimes in hyper-arid deserts.

Our analysis, using an unprecedented sequencing depth for these types of communities, revealed a similar microbial community structure across substrates at the phylum level, which is consistent with previous studies of endolithic communities from multiple deserts (de la Torre *et al.*, 2003; Walker *et al.*, 2005; Dong *et al.*, 2007; Pointing *et al.*, 2007; Azúa-Bustos *et al.*, 2011; DiRuggiero *et al.*, 2013; Li *et al.*, 2013; Crits-Christoph *et al.*, 2016; Armstrong *et al.*, 2016; Lee *et al.*, 2016; Archer *et al.*, 2017; Lacap-Bugler *et al.*, 2017) and suggests that these phyla constitute an ubiquitous “metacommunity” available for the colonization of lithobiontic substrates (Walker and Pace 2007). Indeed, the most abundant taxa at the OTU level all belonged to genera recognized for their resistance to desiccation, radiation, and oligotrophic conditions found in hyper-arid deserts (Friedmann and Ocampo-Friedmann 1995; Potts 1999; Billi *et al.*, 2000; Bull 2011; Krisko and Radman 2013; Mohammadipanah and Wink 2016; Lebre *et al.*, 2017). Our data support a pivotal role for *Cyanobacteria* as primary producers in these endolithic communities as well as potential differences in ecosystem functioning based on the variations in *Cyanobacteria* relative abundance between substrates (Cary *et al.*, 2010; Valverde *et al.*, 2015). As previously proposed, a higher relative abundance of photoautotrophs in the driest environments might indicate reduced capacity for primary production, due to the scarcity of water, hence the decrease in consumers’ relative abundance (Robinson *et al.*, 2015; DiRuggiero *et al.*, 2013). Indeed, salt deliquescence in halite nodules, and therefore conditions conducive to photosynthesis, was continuous in the Salar Grande area of the Atacama Desert while the interior of halite nodules in the driest Yungay area remained wet for only 5362h per year (Robinson *et al.*, 2015; Wierzchos *et al.*, 2012b), matching a higher relative abundance of phototrophs in the Yungay community (Robinson *et al.*, 2015). Here we found low community diversity and abundant *Cyanobacteria* in the granite and ignimbrite rocks, suggesting a harsher environment in these substrates, despite similar inputs of atmospheric water, than in the calcite and gypsum, where the communities were more diverse with a lower abundance of *Cyanobacteria*.

While phylum-level taxa dominated in the four types of substrates, patterns of OTUs’ relative abundance from *Cyanobacteria* and *Actinobacteria* demonstrated preferential association of OTUs with the type of substrate. This partitioning at the OTU phylogenetic level was not recapitulated at higher taxonomic ranks suggesting differential adaptation of close relatives to the substrate’s physical and chemical properties. Substrate-dependent patterns of community structure have been reported for hypolithic quartz and endolithic sandstone communities from the Dry Valleys, Antarctica (Pointing *et al.*, 2009) and for sandstone communities from the Colorado Plateau, USA (Lee *et al.*, 2016). In all cases, while the hypolithic communities were recruited from the surrounding soils, habitat filtering shaped the assembly of the rock communities (Pointing *et al.*, 2009; Makhalanyane *et al.*, 2013; Lee *et al.*, 2016). Community assembly controlled by the substrate was also observed for the hypolithic colonization of meteorites found in Nullarbor Plain, Australia (Tait *et al.*, 2017). While not in desert, partitioning of *Cyanobacteria* was reported for microbial communities in dolomite and limestone substrates, demonstrating mineral preferences of euendolithic communities at the OTU phylogenetic level (Couradeau *et al.*, 2017). Using a sampling strategy that minimized the effect of biogeography and atmospheric water input, our large-scale analysis of four different lithic substrates further provides strong evidence for fine scale diversification of endolithic communities, selectively recruited from a microbial reservoir.

The capacity of a lithic substrate to harbor life, *i.e.* its bioreceptivity, is linked to its chemical, mineralogical and physical properties that provide and maintain sufficient stability and favorable conditions inside the substrate (Walker and Pace 2007; Wierzchos *et al.*, 2012a). Although mineral composition was found to be significantly different between substrates, and suggested a more complex nutrient supply in granite and ignimbrite rocks, low amounts of water soluble ions were detected independently of the substrates’ mineralogy. While in low amounts, nitrogen, phosphate, and other trace elements might not be limiting because of the very low growth rates reported for endolithic communities (Friedmann and Kibler 1980; Ziolkowski *et al.*, 2013; Finstad *et al.*, 2016). Additionally, nitrate levels are typically high in Atacama Desert because of substantial atmospheric deposition (Friedmann and Kibler 1980; Michalski *et al.*, 2004). However, iron limitation would be consistent with the enrichment of genes for iron acquisition and siderophores previously described for ignimbrite and calcite substrates collected in the Atacama Desert and suggesting iron starvation (Crits-Christoph *et al.*, 2016). Unlike previous studies reporting microbe-mineral interactions in specific endolithic habitats (de los Ríos *et al.*, 2004; de los Ríos *et al.*, 2007; Jones and Bennett 2014; Couradeau *et al.*, 2017), we did not find visual evidence for interactions between microorganisms (mostly *Cyanobacteria*) and the surrounding minerals. Neither did we find preferential adhesion of microbial cells to specific minerals. We also did not find OTUs belonging to potential chemolithoautotrophs or anoxygenic photosynthesizers, which metabolisms can modify the pH of the microenvironment, resulting in mineral alterations (Omelon *et al.*, 2007; Jones and Bennett 2014; Jones and Bennett 2017). These findings suggest that rock chemistry in these extreme environments might not be a driving factor for the assembly of endolithic communities.

*Cyanobacteria* photosynthetic activity is essential for endolithic communities and therefore the light intensity and wavelengths reaching the colonization zone within the endolithic habitats could be a limiting factor (Nienow *et al.*, 1988; Boison *et al.*, 2004; Smith *et al.*, 2014; Raanan *et al.*, 2015). Extremely high solar fluxes were measured in the VL and MTQ locations (up to 2,554 µmol photons m^-2^ s^-1^) but rock substrates have the capacity of drastically attenuating incident light and stopping harmful UV radiation (Berner and Evenari 1978; Oren *et al.*, 1995; Matthes *et al.*, 2001; Hughes and Lawley 2003; Horath *et al.*, 2006; Amaral *et al.*, 2007; Hall *et al.*, 2008; Cockell *et al.*, 2010; Cowan *et al.*, 2011; Smith *et al.*, 2014; Wierzchos *et al.*, 2015). Our results clearly showed that differences in light properties of the rock were linked to the location and size of the colonization zone within the substrate but were not related to the richness of the communities or to the relative abundance of phototrophs. For example, the reported transmitted light within the colonization zone of the gypsum at a depth of 2-5 mm was between 0.1 to 1% of the incident PAR (Wierzchos *et al.*, 2015), whereas in granite, narrow fissures between embedded grains of quartz and feldspar known for their translucent properties (Nienow *et al.*, 1988; Schlesinger *et al.*, 2003) allowed a deeper transmission of the light (11% of the incident PAR at a depth of 2 mm, Hall *et al.*, 2008). These values correlate well with our observations of colonization zone’s characteristics but do not link light transmission to community richness. Indeed, the gypsum community was highly diverse with a low abundance of phototrophs and the granite community showed low diversity with a high relative abundance of *Cyanobacteria*. While the community composition of ignimbrite rocks had similar characteristics to that of granite rocks, its dark-colored varnish at the surface and potential light scattering properties from its heterogeneous minerals resulted in a higher absorbance of light by the substrate, restricting the colonization zone to a few mm under its surface (Wierzchos *et al.*, 2013; Crits-Christoph *et al.*, 2016). A large study of limestone rocks of various colors from cliffs of the Niagara Escarpment in Canada similarly reported no significant correlation between light penetration and community richness (Matthes *et al.*, 2001). As such, light is likely the major driver for the localization of the colonization zone within the rock but does not appear to impact community’s richness or *Cyanobacteria’s* abundance. Spectral tuning of photosystem II has been reported in *Cyanobacteria* (Croce and van Amerongen 2014) and experimentally demonstrated with *Synechococcus* variants in microbial mats from hot springs (Becraft *et al.*, 2015), suggesting potential adaptations to low quantum flux densities for endolithic *Cyanobacteria*.

Liquid water is essential for life and required for photosynthetic activity (Palmer and Friedmann 1990; Lange *et al.*, 1993;). While our measurements indicated similar inputs in atmospheric water (rain events and RH) for all sampling locations, we found that the diversity and composition of the communities were highly dependent on the capacity of the substrate to retain liquid water within the lithic habitat: highly diverse communities were associated with substrates that have high potential for water retention (calcite and gypsum) whereas low potential for water retention was linked to low community diversity (granite and ignimbrite). The presence of cracks and fractures and large pores in the gypsum allow water to penetrate easily in the substrate where it is further retained by sepiolite, a magnesium silicate clay with high efficiency for water absorption and retention (Caturla *et al.*, 1999). Indeed, the micro-architectural structure we observed in close proximity of the cells may act as a sponge, enhancing the moisture content within the gypsum rocks (Wierzchos *et al.*, 2015; Vítek *et al.*, 2016). In the calcite habitat, water penetrating cracks and fissures connected to the rock surface will infiltrate deeper within the rocks and be less susceptible to evaporation (DiRuggiero *et al.*, 2013). Additionally, because of the high thermal conductivity properties of calcite, dew can be formed on its surface (DiRuggiero *et al.*, 2013), thus enhancing the water budget for the community (Agam and Berliner 2006; Budel *et al.*, 2008; Kidron and Temina 2013). In contrast, granite is a highly weathered substrate devoid of crust, where water is easily transported in and out of the rock, thus limiting the time it is available to the community. In ignimbrite, small bottle-shaped pores can trap water after short rainfall events, however it has been shown that many pores were unconnected, thus restricting the water budget and making water scarcely accessible to microorganisms (Wierzchos *et al.*, 2013; Cámara *et al.*, 2014).

While the physical structure of the rock, i.e. the size of the cracks, fissures, and pores and their connection to the surface, is an essential factor influencing the water retention capabilities of the rock substrate, the space – size and shape - available for colonization might also be an important factor impacting the habitat microenvironment. For example, in reduced space habitats, such as ignimbrite and granite, the small pores might lead to diffusion limitation of nutrients and metabolic by-products within the community, thus impacting metabolic rates (Johnston and Vestal 1991; Ziolkowski *et al.*, 2013; Chan *et al.*, 2013), community functioning, and ultimately the diversity of the microbial assemblages (Crits-Christoph *et al.*, 2016). In ignimbrite, these diffusional constraints are exacerbated by additional stress to the community from competitions for spare resources and the effects of antimicrobial compounds (Crits-Christoph *et al.*, 2016). In contrast, the large spaces found in gypsum and calcite rocks allow for better diffusion rates and interactions between large aggregates of microorganisms, promoting nutrient exchanges and a diversity of metabolisms (Wierzchos *et al.*, 2015; Crits-Christoph *et al.*, 2016).

In conclusion, we argue that the architecture of the rock (*sensu* Wierzchos *et al.*, 2015), *i.e*. the space available for colonization, embodied by the size of the cracks, fissures, and pores and their connection to the surface is linked to water retention and is ultimately the driver of community diversity and composition in hyper-arid desert endolithic communities. While light is essential to the primary producers of the community, reduce light transmission can be accommodated by spatial variation in colonization zones between substrates. However, further studies of endolithic communities at the functional level and using 3-dimensional microscopic methods might reveal unexpected functional adaptations of community members to their unique habitat.

## EXPERIMENTAL PROCEDURES

The following section is a summary of the Material and Methods used in this study. Detailed procedures are available in the **Supplementary Information**.

### Site description and sampling

Colonized rocks were collected in the Atacama Desert in December 2015 from two locations distant by 100 km: Valle de la Luna area (VL) (GPS coordinates 22°54’S; 068°15’W; 2619 m.a.s.l.) and Monturaqui area (MTQ) (GPS coordinates 23°57’S; 068°10’W; 2868 m.a.s.l) (**Figure 1)**. Both locations are plateaus (photos in **Supplementary Information S1**) located in a N-S trending depression of the Cordon de Lila Range. These areas exhibit a pronounced rain shadow effect by the western slope of the central Andes from 15° to 23°S (DiRuggiero *et al.*, 2013; Wierzchos *et al.*, 2015). Calcite rocks were harvested in VL (**Supplementary Information S1-B1 and B2**). Gypsum (also considered as gypcrete formation) and ignimbrite rocks were harvested in MTQ and were found side by side in the field (**Supplementary Information S1-C1 and C2**). In MTQ area but 25 km to the west, just at the rim of the meteorite impact Monturaqui crater (GPS coordinates 23°55’S; 068°15’W; 3011 m.a.s.l.; Sanchez and Cassidy 1966), very scarce colonized fragments of granite rocks were found sparsely distributed in the field. For each sampling location, rocks were randomly collected within a 50 m^2^ area. All samples were packed in sterile Whirlpack^®^ bags, and stored at room temperature in the dark before further processing.

### Microclimate data

Microclimate data were recorded using an Onset HOBO^®^ Microweather Station Data Logger (H21-001), as previously described (Wierzchos *et al.*, 2015). Air temperature (T), air relative humidity (RH in %) and Photosynthetic Active Radiation (PAR in µmol photons m^-2^s^-1^.) were recorded from May 21^st^, 2013 to December 12^th^, 2015 (33 months) for VL. Microclimate data for MTQ were recorded from January 2011 to February 2013 (22 months) as described previously (Wierzchos *et al.*, 2015). Rainfall data for the both sampling locations were obtained from DiRuggiero *et al.*, (DiRuggiero *et al.*, 2013).

### Electron Microscopy analyses

Colonized rock samples were processed for scanning electron microscopy in back scattered electron mode (SEM-BSE) and/or for energy dispersive X-ray spectroscopy (EDS) microanalysis according to methods by Wierzchos *et al.* (Wierzchos and Ascaso 1994; Wierzchos *et al.*, 2011). Prepared samples (**Supplementary Material and Methods)** were observed using a scanning electron microscope (FEI Quantum 200) equipped with a solid-state, four diodes BSE detector and an auxiliary X-ray EDS microanalytically system (INCA Oxford).

### X-Ray Diffraction analyses

The mineralogical composition of the granite was studied by X-ray powder diffraction (XRD) using a Bruker D8 ADVANCE diffractometer with graphite-monochromated CuK(α) radiation and a linear VANTEC detector. XRD patterns were obtained from powdered samples. Phase identification was performed using the crystallographic database Powder Diffraction File (PDF-2, 1999) from the International Centre for Diffraction Data (ICDD). A semi-quantitative analysis of the phases was performed using the normalized reference intensity ratio (RIR) method (Chung 1974) and the values for each phase given by the powder diffraction database (ICDD).

### Water soluble ions analyses

Major cations and ions were analyzed by Inductively Coupled Plasma-Atomic Emission Spectrometry (EPA-ICP method 200.7) and by Ion Chromatography (EPA – IC method 300.0), respectively, by Inter-Mountain Labs (Sheridan, WY), using 10 grams of fine powder from each substrate, prepared as previously described (Barrett *et al.*, 2009). Duplicate samples were processed for calcite, ignimbrite and gypsum, but not for granite because of the limited number of samples available.

### DNA extraction procedures

At least 4 individual rocks were processed per substrate totalizing 47 samples. The width, maximum and minimum depth of the colonization zone were measured for each substrate before scraping and grounding the colonization zone for DNA extraction. For calcite, ignimbrite and granite, DNA extraction was performed using 0.25g of samples and the PowerSoil kit (MoBio Laboratories Inc., Solana Beach, CA), following the manufacturer’s recommendations. For gypsum samples, we designed a custom protocol by combining sonication, enzymatic lysis and the PowerBiofilm kit (MoBio Laboratories Inc., Solana Beach, CA), with minor modifications (**Supplementary Material and Methods**). All DNA concentrations were measured with the Qubit fluorometer dsDNA HS Assay kit (Life Technologies, Carlsbad, CA). Ignimbrite rocks (n=3) were processed using both protocols to validate the robustness of the DNA isolation procedures and DNA duplicates were processed to validate the robustness of the analytical pipeline (**Supplementary Information S2**).

### 16S rRNA gene libraries preparation and sequencing

A two-step PCR strategy was used to prepare the sequencing libraries, as previously described (Robinson *et al.*, 2015). DNA was amplified using primers 338F and 806R (V3-V4 hypervariable region) barcoded for multiplexing; amplicons from 2 PCR reactions were pooled after the first step. Illumina paired-end sequencing (2 x 250bp) was performed using the MiSeq platform at the Johns Hopkins Genetic Resources Core Facility (GRCF), Libraries from 3 samples were used on all sequencing runs to test for batch effect (**Supplementary Information S2**).

### Computational analysis

After demultiplexing and barcode removal, sequence reads with phred score < 20 and length <100bp were discarded using sickle (Joshi and Fass 2011), representing only 2% of the initial reads count. The Qiime package (v1.6.0) was used to further process the sequences (Caporaso *et al.*, 2010) and diversity metrics were calculated based on OTUs at the 0.03% cutoff against the SILVA database release 123 (Quast *et al.*, 2013). The resulting OTUs table was filtered of the rare OTUs (total abundance across all samples ≤ 0.2%), representing 29% of the initial count (1183 OTUs). Spearman correlations were performed using cor() in R.

The 30% most abundant OTUs for *Cyanobacteria* and *Actinobacteria* were analyzed with R using the pheatmap (Kolde 2015) and DESeq2 packages (Love *et al.*, 2014). Main OTUs were aligned in NCBI GenBank using Clustal W 1.4 software (Thompson *et al.*, 1994). Phylogenetic trees were constructed in MEGA 6.0 using the Maximum Likelihood method (Tamura *et al.*, 2011) and the Kimura 2-parameter model (Kimura 1980). Trees were finally visualized and annotated using the iTOL tool (Letunic and Bork 2016).

### Data submission

The datasets supporting the results of this article are available National Centre for Biotechnology Information under BioSample numbers SAMN07500207-253 and SAMN07519406-418, and BioProject ID PRJNA398025.

## AUTHOR CONTRIBUTIONS

VM and JDR designed and performed the research and wrote the manuscript. JDR and JW conceived the original project. JW and CA performed the microscopy; OA performed the chemical and mineralogy analyses; MCC and MD contributed to the molecular data and analysis; JDR, VM, JW, CA, OA, and MMC participated in sampling and microclimate data acquisition. All authors contributed to editing and revising the manuscript and approved this version for submission.

## ACKNOWLEDGMENTS

This work was founded by grants NNX15AP18G and NNX15AK57G from NASA, grant DEB1556574 from the National Science Foundation to JDR and grant CGL2013-42509P from the Spanish Ministry of Economy, Industry and Competitiveness to JW, CA, OA, JDR and MMC. The MNCN-CSIC, Madrid, Spain is acknowledged for microscopy services.

## REFERENCES

Agam, N., and Berliner, P.R. (2006) Dew formation and water vapor adsorption in semi-arid environments - A review. J. Arid Environ 65: 572–590.

Amaral, G., Martinez-Frias, J., and Vazquez, L. (2007) UV Shielding Properties of Jarosite vs Gypsum: Astrobiological Implications for Mars. World App Sci J 2:112–116.

Archer, S.D.J. et al. (2017) Endolithic microbial diversity in sandstone and granite from the McMurdo Dry Valleys, Antarctica. Polar Biol 40: 997–1006.

Armstrong, A. et al. (2016) Temporal dynamics of hot desert microbial communities reveal structural and functional responses to water input. Scientific Reports 29:34434.

Ascaso, C. and Wierzchos, J. (2002) New approaches to the study of Antarctic lithobiontic microorganisms and their inorganic traces, and their application in the detection of life in Martian rocks. International Microbiol 5:215–222.

Azúa-Bustos, A. et al. (2011) Hypolithic Cyanobacteria Supported Mainly by Fog in the Coastal Range of the Atacama Desert. Microb Ecol 61:568–581.

Barnett, T.P., Adam, J.C. and Lettenmaier, D.P. (2005) Potential impacts of a warming climate on water availability in snow-dominated regions. Nature 438:303–9.

Barrett, J., Ball, B. and Simmons, B. (2009) Standard Procedures for Soil Research in the McMurdo Dry Valleys LTER. pp.1–32.

Becraft, E.D. et al. (2015) The molecular dimension of microbial species: 1. Ecological distinctions among, and homogeneity within, putative ecotypes of Synechococcus inhabiting the cyanobacterial mat of Mushroom Spring, Yellowstone National Park. Front Microbiol 6:1–18.

Berner, T. and Evenari, M. (1978) The Influence of Temperature and Light Penetration on the Abundance of the Hypolithic Algae in the Negev Desert of Israel. Oecologia 33:255–260.

Billi, D. et al. (2000) Ionizing-radiation resistance in the desiccation-tolerant cyanobacterium Chroococcidiopsis. Appl Environ Microbiol 66:1489–1492.

Boison, G. et al. (2004) Bacterial Life and Dinitrogen Fixation at Gysum Rock. Appl Environ Microbiol 70:7070–7077.

Büdel, B. et al. (2008) Dewfall as a water source frequently activates the endolithic cyanobacterial communities in the granites of Taylo Valley, Antarctica. J Phycol 44:1415–1424.

Bull, A.T. (2011) Actinobacteria of the extremobiosphere. In Extremophiles Handbook. Koki Horikoshi (ed.). Springer Japan, pp. 1204–1231.

Cámara, B. et al. (2014) Ignimbrite textural properties as determinants of endolithic colonization patterns from hyper-arid Atacama Desert. Int Microbiol 17:235–247.

Canfora, L. et al. (2016) Compartmentalization of gypsum and halite associated with cyanobacteria in saline soil crusts. FEMS Microb Ecol 92:1–13.

Caporaso, J.G. et al. (2010) QIIME allows analysis of high-throughput community sequencing data. Nature Methods 7:335–336.

Cary, S.C. et al. (2010) On the rocks: the microbiology of Antarctic Dry Valley soils. Nature reviews. Microbiol 8:129–138.

Caturla, F., Molina-Sabio, M. and Rodriguez-Reinoso, F. (1999) Adsorption-desorption of water vapor by natural and heat-treated sepiolite in ambient air. Appl Clay Sci 15:367–380.

Chan, Y. et al. (2013) Functional ecology of an Antarctic Dry Valley. Proc Natl Acad Sci U S A, 110:8990–8995.

Chung, F.H. (1974) Quantitative interpretation of X-ray diffraction patterns of mixtures. II. Adiabatic principle of X-ray diffraction analysis of mixtures. J Appl Crystal 7:526–531.

Cockell, C., McKay, C.P. and Omelon, C. (2002) Polar endoliths – an anti-correlation of climatic extremes and microbial biodiversity. Internl J Astrobiol 1:305–310.

Cockell, C.S. et al. (2010) The microbe-mineral environment and gypsum neogenesis in a weathered polar evaporite. Geobiol 8:293–308.

Cordero, R.R. et al. (2014) The world’s highest levels of surface UV. Photochem Photobiol Sci 13:70–81.

Couradeau, E. et al. (2017) Diversity and mineral substrate preference in endolithic microbial communities from marine intertidal outcrops (Isla de Mona, Puerto Rico). Biogeosci 14:311–324.

Cowan, D.A. et al. (2011) Distribution and abiotic influences on hypolithic microbial communities in an Antarctic Dry Valley. Polar Biol 34:307–311.

Crits-Christoph, A. et al. (2016) Phylogenetic and Functional Substrate Specificity for Endolithic Microbial Communities in Hyper-Arid Environments. Front Microbiol 7:1–15.

Croce, R. and van Amerongen, H. (2014) Natural strategies for photosynthetic light harvesting. Nature Chem Biol 10:492–501.

Davila, A.F. et al. (2008) Facilitation of endolithic microbial survival in the hyperarid core of the Atacama Desert by mineral deliquescence. J Geophys Res: Biogeosci 113:1–9.

de la Torre, J. et al. (2003) Microbial diversity of cryptoendolithic communities from the McMurdo Dry Valleys, Antarctica. Appl Environ Microbiol 69:3858–3867.

de los Ríos, A. et al. (2010) Comparative analysis of the microbial communities inhabiting halite evaporates of the Atacama Desert. Intern Microbiol 13:79–89.

de los Ríos, A. et al. (2005) Ecology of endolithic lichens colonizing granite in continental Antarctica. The Lichenologist 37:383.

de los Ríos, A. et al. (2004) Exploring the physiological state of continental Antarctic endolithic microorganisms by microscopy. FEMS Microbiol Ecol 50:143–152.

de los Ríos, A. et al. (2007) Ultrastructural and genetic characteristics of endolithic cyanobacterial biofilms colonizing Antarctic granite rocks. FEMS Microbiol Ecol 59:386–395.

DiRuggiero, J. et al. (2013) Microbial colonisation of chasmoendolithic habitats in the hyper-arid zone of the Atacama Desert. Biogeosci 10:2439–2450.

Dong, H. et al (2007) Endolithic Cyanobacteria in soil gypsum: Occurences in Atacama (Chile), Mojave (United States), and Al-Jafr Basin (Jordan) Deserts. J Geophys Res: Biogeosci 112:1–11.

Finstad, K., Pfeiffer M., McNicol G., Barnes J., Demergasso C., Chong G. and Amundson R. (2016) Rates and geochemical processes of soil and salt crust formation in Salars of the Atacama Desert, Chile. Geoderma 284:57–72.

Friedmann, E.I. (1980) Endolithic Microbial Life in Hot and Cold Deserts. Origins Of Life, 10:223–235.

Friedmann, E.I., Hua, M. and Ocampo-Friedmann, R. (1988) Cryptoendolithic lichen and cyanobacterial communities of the Ross Desert, Antarctica. Polarforschung 58:251–259.

Friedmann, E.I. and Kibler, A.P. (1980) Nitrogen economy of endolithic microbial communities in hot and cold deserts. Microb Ecol 6:95–108.

Friedmann, I.E. and Ocampo-Friedmann, R. (1995) A primitive cyanobacterium as pioneer microorganism for terraforming Mars. Adv Space Res 15:243–246.

Golubic, S., Friedmann, I. and Schneider, J. (1981) The lithobiontic ecological niche, with special reference to microorganisms. J Sedimentary Res 51:475–478.

Hall, K., Guglielmin, M. and Strini, A. (2008) Weathering of granite in Antarctica: I. Light penetration into rock and implications for rock weathering and endolithic communities. Earth Surf Process Landforms 33:295–307.

Horath, T., Neu, T.R. and Bachofen, R. (2006) An endolithic microbial community in dolomite rock in Central Switzerland: Characterization by reflection spectroscopy, pigment analyses, scanning electron microscopy, and laser scanning microscopy. Microb Ecol 51:353–364.

Hughes, K.A. and Lawley, B. (2003) A novel Antarctic microbial endolithic community within gypsum crusts. Environ Microbiol 5:555–565.

Johnston, C.G. and Vestal, J.R. (1991) Photosynthetic carbon incorporation and turnover in Antarctic cryptoendolithic microbial communities: Are they the slowest-growing communities on earth? Appl EnvironMicrobiol 57:2308–2311.

Jones, A.A. and Bennett, P.C. (2017) Mineral ecology: Surface specific colonization and geochemical drivers of biofilm accumulation, composition, and phylogeny. Front Microbiol 8:1–14.

Jones, A.A. and Bennett, P.C. (2014) Mineral Microniches Control the Diversity of Subsurface Microbial Populations. Geomicrobiol J 31:246–261.

Joshi, N. and Fass, J. (2011) Sickle: A sliding-window, adaptive, quality-based trimming tool for FastQ files (version 1.33) [Software]. Available at https://github.com/najoshi/sickle.

Kidron, G.J. and Temina, M. (2017) Non-rainfall water input determines lichen and cyanobacteria zonation on limestone bedrock in the Negev Highlands. Flora 229:71–79.

Kidron, G.J. and Temina, M. (2013) The Effect of Dew and Fog on Lithic Lichens Along an Altitudinal Gradient in the Negev Desert. Geomicrobiol J 30:281–290.

Kimura, M. (1980) A simple method for estimating evolutionary rates of base substitutions through comparative studies of nucleotide sequences. J Molr Evol 16:111–120.

Kolde, R. (2015) pheatmap: Pretty Heatmaps. *R package version 1.0.8*. http://CRAN.R-project.org/package=pheatmap.

Krisko, A. and Radman, M. (2013) Biology of Extreme Radiation Resistance: the way of Deinococcus radiodurans. Cold Spring Harb Perspect Biol 5:1–12.

Lacap-Bugler, D.C. et al. (2017) Global Diversity of Desert Hypolithic Cyanobacteria. Front Microbiol 8:1–13.

Lange, O.L. et al. (1993) Further Evidence That Activation of Photosyntesis By Dry Cyanobacterial Lichens Requires Liquid Water. Lichenologist 25:175–189.

Lebre, P.H., De Maayer, P. and Cowan, D.A. (2017) Xerotolerant bacteria: surviving through a dry spell. Nature Rev Microbiol 15:285–296.

Lee, K.C. et al. (2016) Niche filtering of bacteria in soil and rock habitats of the Colorado Plateau Desert, Utah, USA. Front Microbiol 7:1–7.

Letunic, I. and Bork, P. (2016) Interactive tree of life (iTOL) v3: an online tool for the display and annotation of phylogenetic and other trees. Nucl acids Res 44:W242–5.

Li, S. et al. (2013) Phylogenetic diversity of endolithic bacteria in Bole granite rock in Xinjiang. Acta Ecol Sinica 33:178–184.

Love, M.I., Huber, W. and Anders, S. (2014) Moderated estimation of fold change and dispersion for RNA-seq data with DESeq2. Genome Biol 15:550.

Makhalanyane, T.P. et al. (2013) Evidence of species recruitment and development of hot desert hypolithic communities. Environ Microbioly Rep 5:219–224.

Makhalanyane, T.P. et al. (2015) Microbial ecology of hot desert edaphic systems. FEMS Microbiol Rev 39:203–221.

Matthes, U., Turner, S.J. and Larson, D.W. (2001) Light Attenuation by Limestone Rock and Its Constraint on the Depth Distribution of Endolithic Algae and Cyanobacteria. Int J Plant Sci 162:263–270.

McKay, C.P. and Friedmann, E.I. (1985) The cryptoendolithic microbial environment in the Antarctic cold desert: Temperature variations in nature. Polar Biol 4:19–25.

Michalski, G., Böhlke, J.K. and Thiemens, M. (2004) Long term atmospheric deposition as the source of nitrate and other salts in the Atacama Desert, Chile: New evidence from mass-independent oxygen isotopic compositions. Geochim Cosmochim Acta 68:4023–4038.

Mohammadipanah, F. and Wink, J. (2016) Actinobacteria from arid and desert habitats: Diversity and biological activity. Front Microbiol 6:1–10.

Nienow, J.A., McKay, C. and Friedmann, E.I. (1988) The Cryptoendolithic Microbial Environment in the Ross Desert of Antarctica: Light in the Photosynthetically Active Region. Microb Ecol 16:271–289.

Omelon, C.R. (2008) Endolithic Microbial Communities in Polar Desert Habitats. Geomicrobiol J 25:404–414.

Omelon, C.R., Pollard, W.H. and Ferris, F.G. (2006) Environmental controls on microbial colonization of high Arctic cryptoendolithic habitats. Polar Biol 30:19–29.

Omelon, C.R., Pollard, W.H. and Ferris, F.G. (2007) Inorganic species distribution and microbial diversity within high arctic cryptoendolithic habitats. Microb Ecol 54:740–752.

Oren, A., Kuhl, M. and Karsten, U. (1995) An endoevaporitic microbial mat within a gypsum crust: zonation of phototrophs, photopigments, and light penetration. Mar Ecol Prog Ser 128:151–159.

Palmer and Friedmann (1990) Water relations, thallues structure and photosynthesis in Negev Desert lichens. New Phytol 116:597–603.

Pointing, S.B. et al. (2009) Highly specialized microbial diversity in hyper-arid polar desert. Proc Nat Acad Sci U S A 106:19964–19969.

Pointing, S.B. et al. (2007) Hypolithic community shifts occur as a result of liquid water availability along environmental gradients in China’s hot and cold hyperarid deserts. Environ Microbiol 9:414–424.

Pointing, S.B. and Belnap, J. (2012) Microbial colonization and controls in dryland systems. Nature Rev Microbiol 10:654.

Potts, M. (1999) Mechanisms of desiccation tolerance in Cyanobacteria. Europ J Phycol 34:319–328.

Quast, C. et al. (2013) The SILVA ribosomal RNA gene database project: Improved data processing and web-based tools. Nucl Acids Res 41:590–596.

Raanan, H. et al. (2015) Three-dimensional structure and cyanobacterial activity within a desert biological soil crust. Environ Microbiol 18:372–383.

Robinson, C.K. et al. (2015) Microbial diversity and the presence of algae in halite endolithic communities are correlated to atmospheric moisture in the hyper-arid zone of the Atacama Desert. Environ Microbiol 17:299–315.

Sanchez, J. and Cassidy, W. (1966) A previously undescribed meteorite crater in Chile. J Geophys Res 71:4891–4895.

Schlesinger, W.H. et al. (2003) Community composition and photosynthesis by photoautotrophs under quartz pebbles, southern Mojave Desert. Ecology 84:3222–3231.

Smith, H.D. et al. (2014) Comparative analysis of cyanobacteria inhabiting rocks with different light transmittance in the Mojave Desert: a Mars terrestrial analogue. Int J Astrobiol 13:1–7.

Stivaletta, N. et al. (2010) Biomarkers of Endolithic Communities within Gypsum Crusts (Southern Tunisia). Geomicrobiol J 27:101–110.

Stomeo, F. et al. (2013) Hypolithic and soil microbial community assembly along an aridity gradient in the Namib Desert. Extremophiles 17:329–337.

Tait, A.W. et al. (2017) Microbial Populations of Stony Meteorites: Substrate Controls on First Colonizers. Front Microbiol 8:1–14.

Tamura, K. et al. (2011) MEGA5: Molecular evolutionary genetics analysis using maximum likelihood, evolutionary distance, and maximum parsimony methods. Mol Biol Evol 28:2731–2739.

Tang, Y. et al. (2012) Endolithic bacterial communities in Dolomite and Limestone rocks from the Nanjiang Canyon in Guizhou Karst Area (China). Geomicrobiol J 29:213–225.

Tang, Y., Cheng, J. and Lian, B. (2016) Characterization of Endolithic Culturable Microbial Communities in Carbonate Rocks from a Typical Karst Canyon in Guizhou (China). Pol J Microbiol 65:413–423.

Thompson, J.D., Higgins, D.G. and Gibson, T.J. (1994) CLUSTAL W: Improving the sensitivity of progressive multiple sequence alignment through sequence weighting, position-specific gap penalties and weight matrix choice. Nucl Acids Res 22:4673–4680.

Valverde, A. et al. (2015) Cyanobacteria drive community composition and functionality in rock-soil interface communities. Mol Ecol 24:812–821.

Vítek, P. et al. (2016) Raman imaging in geomicrobiology: endolithic phototrophic microorganisms in gypsum from the extreme sun irradiation area in the Atacama Desert. Anal Bioana Chem 408:4083–4092.

Walker, J., Spear, J. and Pace, N. (2005) Geobiology of a microbial endolithic community in the Yellowstone geothermal environment. Nature 434:1011–1014.

Walker and Pace, N. (2007) Endolithic Microbial Ecosystems. Ann Rev Microbiol 61:331–347.

Wierzchos et al. (2013) Ignimbrite as a substrate for endolithic life in the hyper-arid Atacama Desert: Implications for the search for life on Mars. Icarus 224:334–346.

Wierzchos et al. (2012b) Novel water source for endolithic life in the hyperarid core of the Atacama Desert. Biogeosci 9:2275–2286.

Wierzchos, J. et al. (2015) Adaptation strategies of endolithic chlorophototrophs to survive the hyperarid and extreme solar radiation environment of the Atacama Desert. Front Microbiol 6:1–11.

Wierzchos, J. et al. (2011) Microbial colonization of Ca-sulfate crusts in the hyperarid core of the Atacama Desert: Implications for the search for life on Mars. Geobiol 9:.44–60.

Wierzchos, J. and Ascaso, C. (1994) Application of back**‐**scattered electron imaging to the study of the lichen-rock interface. J Microscopy 175:54–59.

Wierzchos, J. and Ascaso, C. (2001) Life, decay and fossilisation of endolithic microorganisms from the Ross Desert, Antarctica. Polar Biol 24:863–868.

Wierzchos, J., Ascaso, C. and McKay, C.P. (2006) Endolithic cyanobacteria in halite rocks from the hyperarid core of the Atacama Desert. Astrobiol 6:415–422.

Wierzchos, J., de los Ríos, A. and Ascaso, C. (2012a) Microorganisms in desert rocks: The edge of life on Earth. Intl Microbiol 15:173–183.

Ziolkowski, L.A. et al. (2013) Radiocarbon Evidence of Active Endolithic Microbial Communities in the Hyperarid Core of the Atacama Desert. Astrobiol 13:607–616.

